# ClusterMap: Comparing analyses across multiple Single Cell RNA-Seq profiles

**DOI:** 10.1101/331330

**Authors:** Xin Gao, Deqing Hu, Madelaine Gogol, Hua Li

## Abstract

Single cell RNA-Seq facilitates the characterization of cell type heterogeneity and developmental processes. Further study of single cell profiles across different conditions enables the understanding of biological processes and underlying mechanisms at the sub-population level. However, developing proper methodology to compare multiple scRNA-Seq datasets remains challenging. We have developed ClusterMap, a systematic method and workflow to facilitate the comparison of scRNA profiles across distinct biological contexts. Using hierarchical clustering of the marker genes of each sub-group, ClusterMap matches the sub-types of cells across different samples and provides “*similarity*” as a metric to quantify the quality of the match. We introduce a purity tree cut method designed specifically for this matching problem. We use Circos plot and regrouping method to visualize the results concisely. Furthermore, we propose a new metric “*separability*” to summarize sub-population changes among all sample pairs. In three case studies, we demonstrate that ClusterMap has the ability to offer us further insight into the different molecular mechanisms of cellular sub-populations across different conditions. ClusterMap is implemented in R and available at https://github.com/xgaoo/ClusterMap.

## Introduction

Single cell RNA sequencing (scRNA-Seq) is an advanced technology that sheds light on the high-resolution heterogeneity and dynamics of the transcriptome. Following massive studies on cell sub-types under static conditions, researchers are starting to pursue high-resolution mechanisms of different biological processes. With more complicated experimental designs that include different treatment conditions, different developmental time points, or different tissue types, researchers seek information on heterogeneous changes of sub-populations of cells.

There are many existing methods and packages for identification of cellular sub-types, developmental trajectory, and differential expression analysis in single cell expression analysis (1–10). However, most of them are either restricted to one single cell dataset or focused on batch effect correction across datasets (11, 12). Methods for directly comparing multiple scRNA-Seq datasets across biological conditions to study heterogeneity of changes of sub cell types are still under development. The current common strategy is to combine multiple datasets and then analyze it as a single dataset, which might miss critical information at the cell sub-type level. Some recently published methods attempt to address similar issues (13, 14). Butler et al (13) attempt to align scRNA-seq datasets using canonical-correlation analysis (CCA) so that shared subpopulations across datasets can be compared directly. Scmap (14) maps a single scRNA-seq sample to the reference dataset to annotate the new dataset. Scmap maps each cell to the cluster centroid or the nearest cell in the reference. Here we present ClusterMap, a tool to analyze and compare multiple single cell expression datasets at the cluster level. We focus on the difference across multiple samples under different treatments or conditions. ClusterMap uses marker genes of each sub-group as features for the comparison. By clustering sub-groups and using purity tree cut methods, ClusterMap enables matching and comparison of sub cell types across multiple samples. It also provides the “*similarity*” measurement on matched groups, to assess the confidence of the matching, and “*separability*” to quantify changes within each sub-group across samples. ClusterMap automates systematic comparison analysis through the following steps.

First, we attempt to match the most similar sub-groups across samples. The typical approach is to check the expression of one or a few known marker genes and use that to match the corresponding sub-groups (15–17). This manual step biases the matching towards several pre-selected genes and might not be accurate. ClusterMap provides an objective method to match sub-groups, based on the marker genes of each sub-group identified in each individual dataset. By clustering binary expression patterns of marker genes, we build a hierarchical tree showing distance between sub-groups. This allows sub-group matching even in the absence of known marker genes. Next, we propose a new tree cut method, purity tree cut, to group the most similar sub-groups. This could result in groups matching in a one-to-multiple, multiple-to-multiple, or singleton fashion, which enables the detection of unique or novel sub-types of cells. Meanwhile we introduce *similarity* to quantify the absolute level of the quality of the match, which indicates the percentage of marker genes that overlap between matched groups. Finally, we quantify the changes within each sub-group of cells across samples. These differences can be population size changes, expression changes, or both. We define *separability* to characterize the conditional effect for each sub population, which enables quick and unbiased identification of the most highly affected cell sub-types among all the sample comparisons. ClusterMap also allows visualization of the matched pairs, the *similarity* of pairs, and the cell percentage difference in a Circos plot. The dimensional reduction (t-SNE) plots are then re-colored to display the paired groups.

Overall, ClusterMap provides an easy-to-use and reliable workflow to compare multiple single cell RNA-Seq datasets with complex experimental designs: across various treatments, across time points, and across tissues. We demonstrate the usage and advantages of ClusterMap in three case studies. 1) Analysis of mammary gland epithelial cells across different estrus phases 2) Comparison of immune stimulated PBMCs with control. 3) Comparison of replicates of PBMCs datasets (as a negative control). Our analysis shows precise alignment of the sub-groups in different conditions, accurate characterization of differences between matched sub-groups, and identifaction of unique sub-groups of cells. Our method is a valuable tool for comparison of complex scRNA-seq datasets with multiple treatments or timepoints and offers a deeper understanding of the biological processes at the cellular sub-population level.

## Methods

### Workflow overview

ClusterMap focuses on the analysis of sub-group matching and comparison across single cell RNA-Seq samples. ClusterMap analysis is based on the pre-analysis of each individual dataset. The sub-group definitions, identification of marker genes for each group, and dimension reduction are generated in the pre-analysis step and used as input to ClusterMap. This step can be achieved using the Seurat package (1), 10X Genomics Cell Ranger, or other single cell analysis methods. The summarized workflow of ClusterMap is shown in Figure 1.

**Figure 1.**
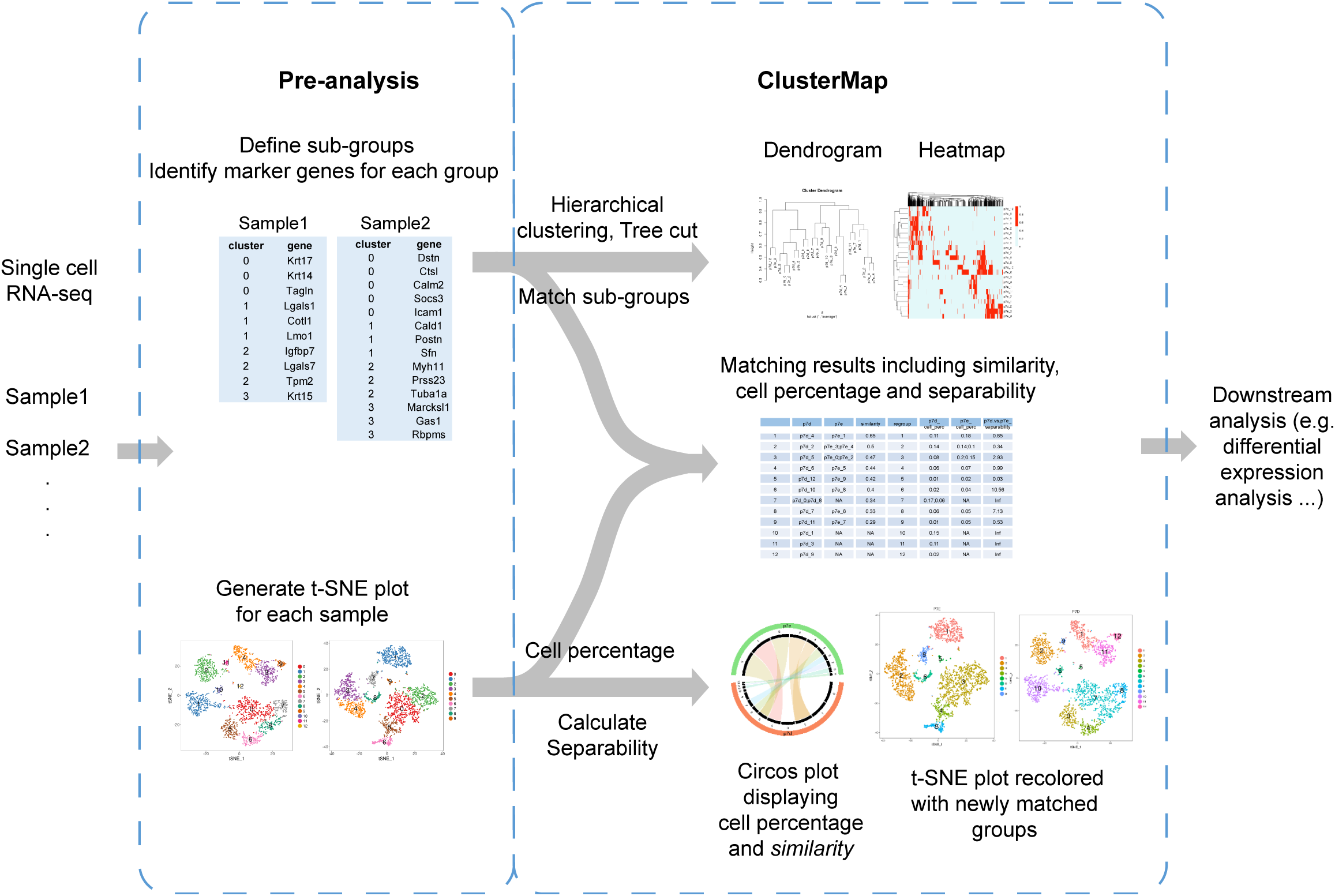
Diagram of the workflow of ClusterMap.

**Figure 2.**
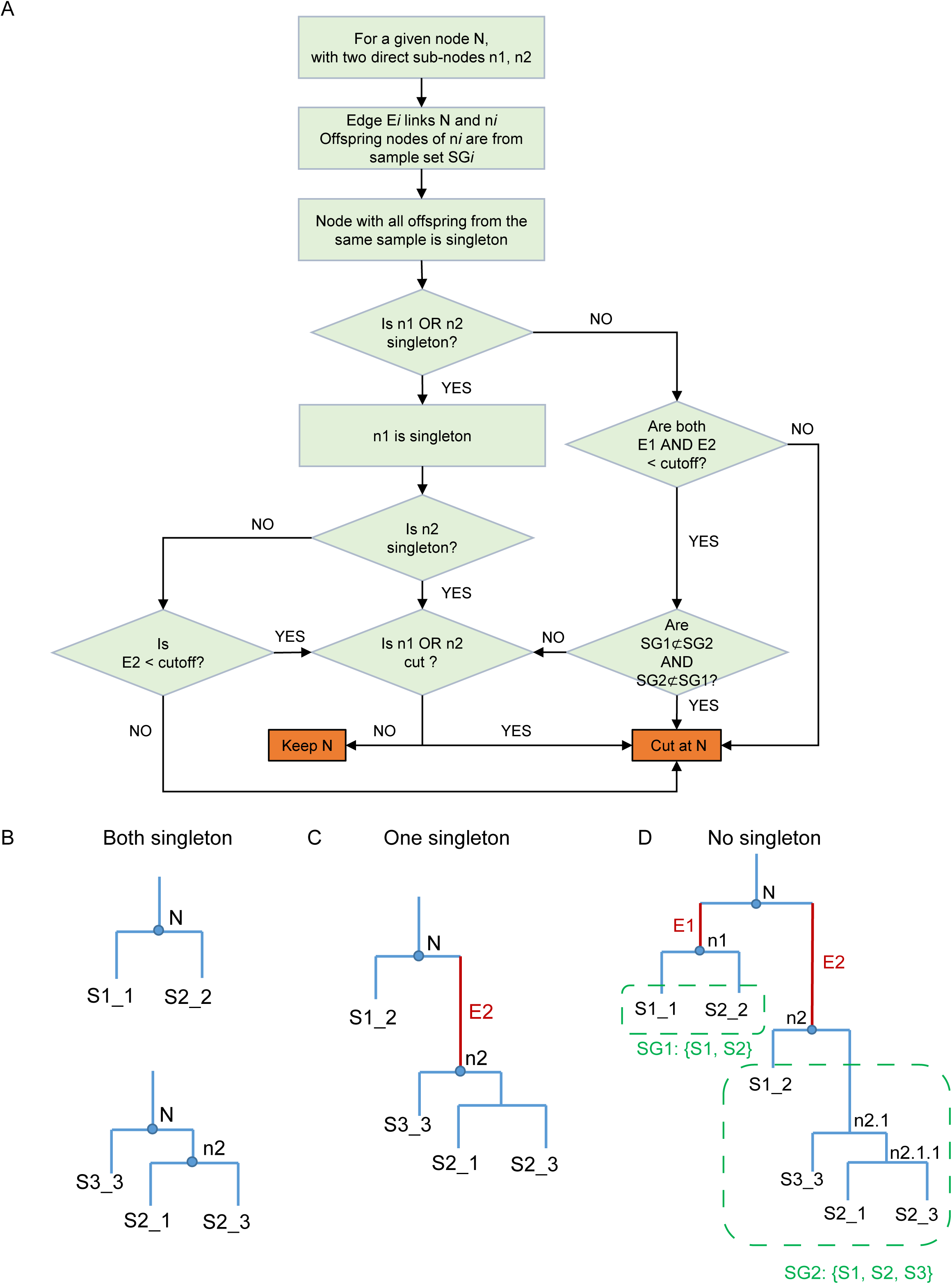
Diagram of the dendrogram tree cut algorithm. A. Purity tree cut algorithm. B-D. Examples of the three conditions in the tree cut. B. Two singletons. C. One singleton. D. No singleton.

ClusterMap uses identified marker genes for each sub-group of each sample as the basic input to match clusters. Through hierarchical clustering analysis of the binary expression patterns of marker genes and the purity tree cut method, the sub-groups identified in the pre-analysis step for each individual dataset will be matched and grouped together with the most similar sub-groups in the other samples. The *similarity* of the matched groups is extracted from the clustering results. With the number of cells (cell percentage) in each sub-group as an additional input, ClusterMap generates a Circos plot to show matched sub-groups between datasets and the compositional changes of the sub-groups across datasets (Figure 1, 3E). The chords link the matched sub-groups, while different chord colors indicate different regroups and the transparency of the chord color indicates the *similarity* of matched groups (more transparent indicates less similar). The widths of the black sectors represent the percentage of the number of cells in each sample. Thus, the sub-population size change is intuited by comparing the sector size of linked groups. New cluster labels will be assigned according to the cluster matching results. If two-dimensional coordinates from a dimensional reduction (t-SNE) plot are provided for each cell, ClusterMap will re-color the plot to coordinate the colors for the matched groups in different samples (Figure 1, 4A-C). This will facilitate the visualization of the matching sub-groups. Finally, ClusterMap will calculate the *separability* to characterize the property changes across samples for each set of matched groups (Figure 1, 3F). *Separability* is calculated for a pair of samples. Pairwise *separability* will be measured if there are more than two samples.

**Figure 3.**
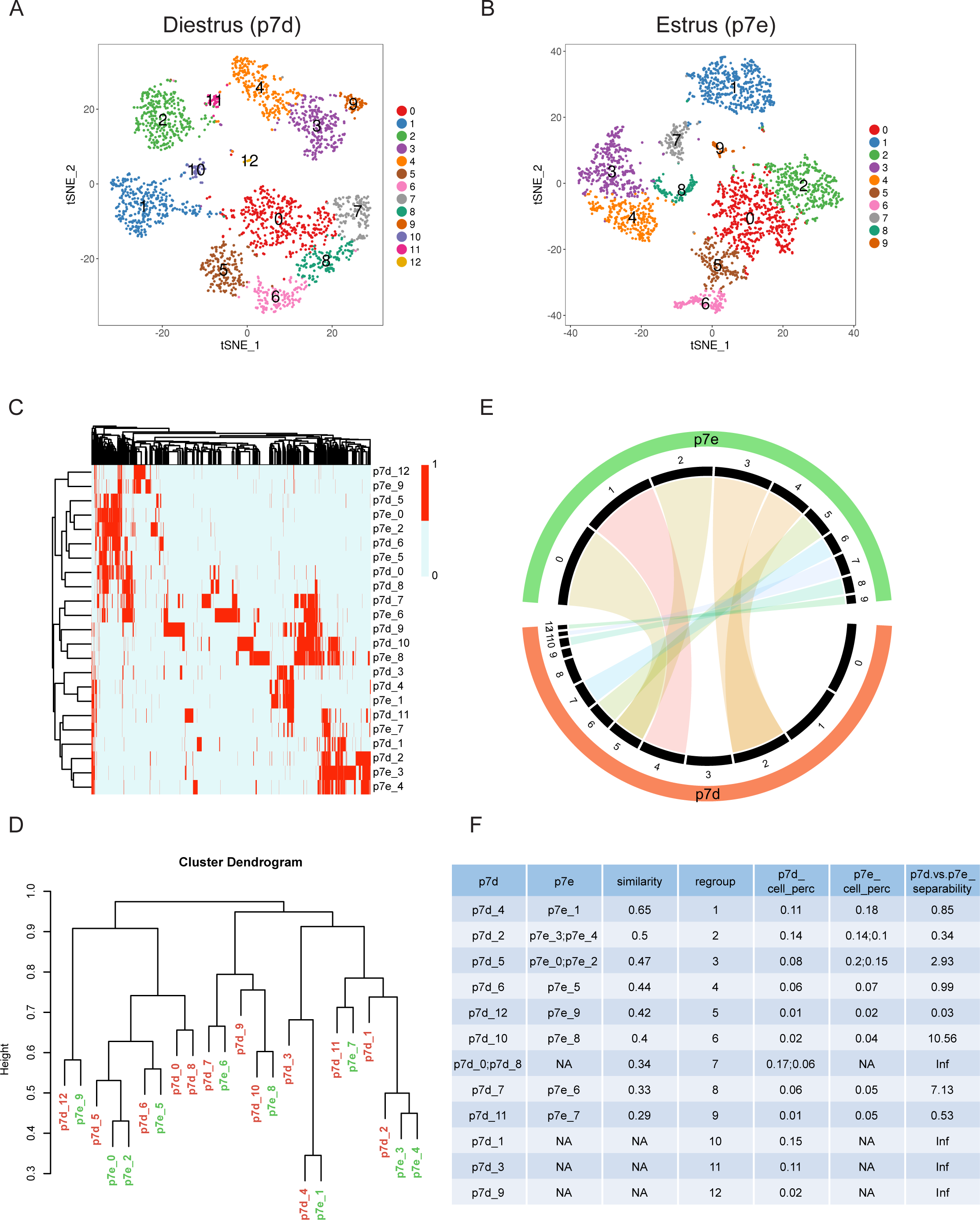
Cluster match of mammary gland epithelial cells at different phases of the estrus cycle. A. Pre-analysis of cells from diestrus phase. B. Pre-analysis of cells from estrus phase. C. Heat map of the hierarchical clustering of subgroups by the existence of marker genes. Each column is a marker gene for one of the subgroups. D. Dendrogram of the hierarchical clustering. E. Circos plot of the matched subgroups. The 13 black sectors highlighted by the red sector represent 13 groups in diestrus as in Figure 3A, while the 10 sectors highlighted by the green sector represent 10 groups in estrus as in Figure 3B. The width of the black sectors represents the percentage of cells in each sample. Matched groups are linked by chord, and the transparency represents the *similarity* of the matched groups, with less transparent indicating more similar. F. ClusterMap results for the quantification of the sample comparisons.

**Figure 4.**
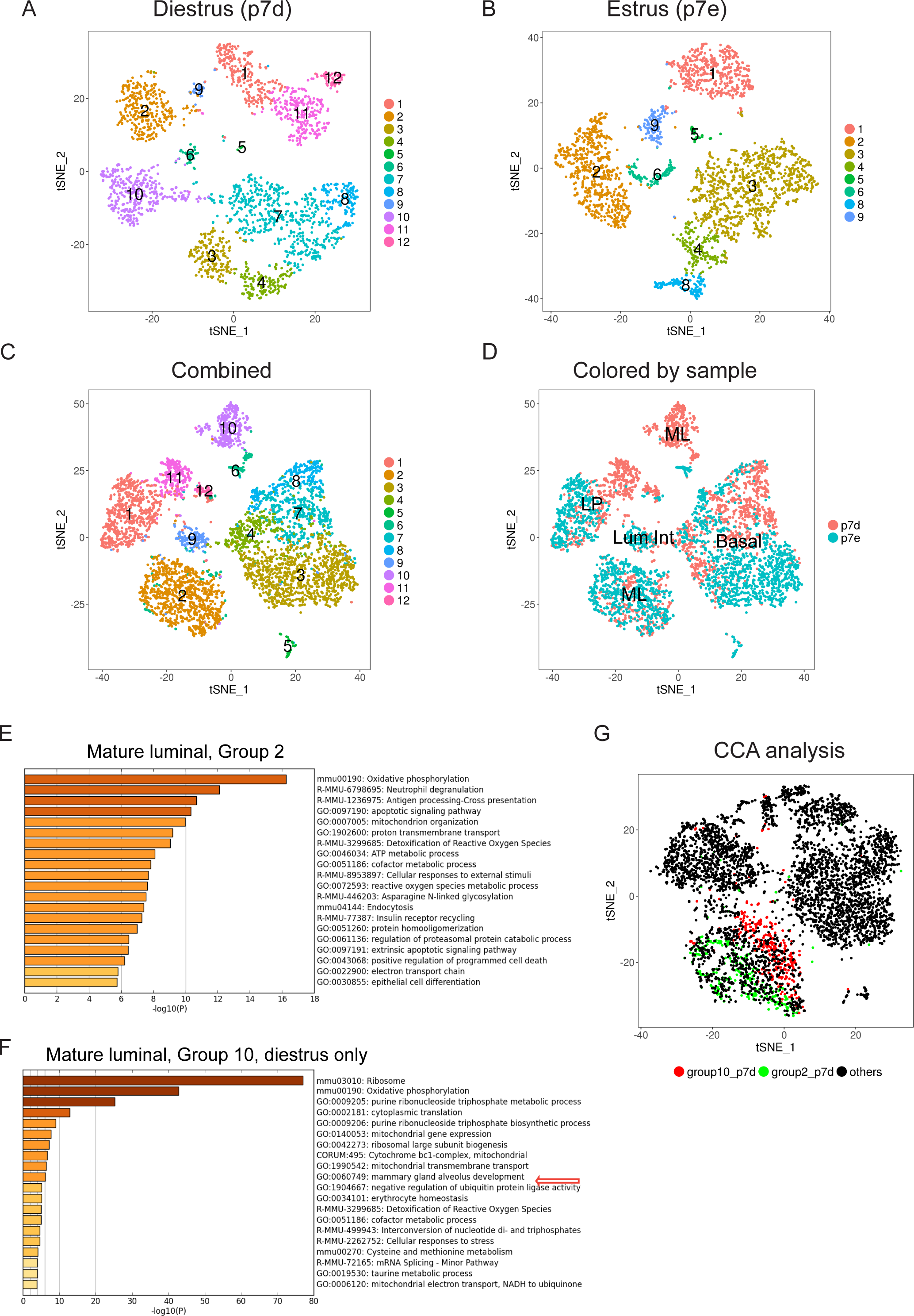
Regrouping of mammary gland epithelial cells. A. Re-colored t-SNE plot based on matching results for diestrus phase. B. Re-colored t-SNE plot based on matching results for estrus phase. C. Re-colored t-SNE plot based on matching results for the combined dataset. Matched groups were recolored in the same color and with the same label through all three t-SNE plots. D. t-SNE plot with cells colored by sample. Based on known markers (Figure S2A-B), we generally assigned group 2, 5, 6, 10 to mature luminal (ML), group 1, 11, 12 to luminal progenitors (LP), group 9 to luminal intermediate (Lum Int), and group 3, 4, 7, 8 to basal cells. E-F. Gene ontology and pathway analysis for the new marker genes of group 2 and 10 in the combined sample (C) using Metascape. G. t-SNE plot of CCA analysis. Cells of group 2 and group 10 in diestrus, defined as in Figure 4A are highlighted in green and red.

Following ClusterMap, differential expression analysis for the most affected group might be a common following step to investigate the difference further. Since many methods have been well established, such as DESeq2, SCDE, BASiCS (9, 18–20), this aspect of analysis is not included in ClusterMap.

### Binary hierarchical clustering

The sub-groups are matched using hierarchical clustering (Figure 1, 3D). The presence (binary) of the marker genes in each group is used to measure the distance between the sub-groups across the samples. The union of all marker genes is used for clustering. We construct a matrix showing expression of all identified marker genes for all sub clusters across multiple datasets. The value of a marker gene is assigned as 1 or 0, depending on whether this gene is identified as the marker gene for a specific sub-cluster or not. Hierarchical clustering of this binary matrix is performed using the average linkage and Jaccard distance. The *similarity* of the matched groups is defined as 1 minus the height of the merging node of the matched groups in the dendrogram, which is the Jaccard index at that node (Figure 1, 3F). It measures the percentage of marker genes that overlap between the matched groups.

Using the binary value of marker genes is superior to using the actual expression level to match sub-groups across samples for a few reasons. First, the expression level of a gene across cells or samples may vary a lot under different conditions. Without proper normalization methods, its direct use will be subject to noise. Binarizing expression tolerates global shifts of the transcriptome, or possible systematic batch effect across datasets. Second, given the high drop-out rate of current scRNA-seq technology, using binary values is more reliable and robust. Consequently, using the Jaccard distance as a measure of *similarity* of sub-groups is more reasonable than Euclidian distance.

### Purity Tree Cut

The purity tree cut algorithm is designed to match the most similar sub-groups from different samples, while avoiding forming large clusters of sub-groups that come from the same sample. Traditional clustering assessment methods, such as the Elbow method, Silhouette index, Dunn index, or other indexes, are not optimal for this purpose, because the origin of samples is disregarded. Our algorithm decomposes the clustering dendrogram from the bottom-up based on both the distance between branches and the purity of the node (Figure 2A). The purity tree cut algorithm decides to keep or trim a given node by checking the following three aspects.

First, we consider the purity of the node. In a dendrogram, the sample set of a node is the set of samples that all its offspring nodes come from. If all the offspring nodes are from the same sample (only one sample is in its sample set), then the node is considered pure, and treated as a singleton node. For example, in Figure 2D, the offspring of node n1 are S1_1 and S2_2. They are from sample S1 and S2. Thus, the sample set of node n1 is {S1, S2} and n1 is not a pure node. For node n2.1.1, the offspring S2_1 and S2_3 are from the same sample S2, thus n2.1.1 is a pure node and is treated as a singleton node (Figure 2D).

Second, we consider the edge length of the two branches. The edge length in a dendrogram is the height difference between the upper node (such as N) and the lower node (such as n1 in Figure 2D). An edge length cutoff controls whether two branches of a node should be merged into one group. If an edge is longer than the cutoff, then one branch is quite different from the other branch, and it will not be merged. The default cutoff is set to 0.1, so less than 10% of the marker genes can be different in order to continue merging two branches.

Third, we consider the overlap between the sub-samples of the two branches.

We search through all the nodes in the dendrogram from the bottom up. For a given node N with two direct sub-nodes n1 and n2, and edges E1 and E2, N will be kept only under one of the three conditions:

1. n1 and n2 are both singleton or pure (Figure 2B).
2. n1 is singleton or pure, AND the edge E2 is shorter than the edge cutoff, AND n2 is not cut based on the sub-nodes of n2 (Figure 2C).
3. Both edge E1 and E2 are shorter than the edge cutoff, AND the two sample sets SG1 and SG2 are not subsets of each other (SG1⊄SG2 AND SG2⊄SG1), AND none of n1 or n2 is trimmed (Figure 2D).

In all other conditions, node N will be removed, and n1 and n2 will form two different groups. Using Figure 2D as an example, based on condition 1, node n1 will be kept, because both S1_1 and S2_2 are singletons. So S1_1 and S2_2 are matched and form a new group Node n2.1.1 is equivalent to a singleton, thus node n2.1 will be kept with two singleton sub-nodes based on condition 1. S2_1, S2_3, and S3_3 will form one matched group. Due to condition 2, node n2 will be trimmed if the edge between n2 and n2.1 is longer than the cutoff, and S1_2 will not be grouped together with the other branch. Node N violates condition 3 and will be trimmed, because SG1 is a subset of SG2.

We generated a random tree with four samples and 10 sub-groups in each sample to test the purity tree cut algorithm (Figure S1A). The results were as expected, similar sub-groups are grouped together but avoid forming big groups from the same sample (Figure S1B). If we increase the edge cutoff, more sub-groups merge into bigger groups but with lower *similarity* (Figure S1C). The following case studies also suggest that the tree cutting results match the underlying biological expectations.

The purity tree cut algorithm will match a group with its most closely related sub-group from another sample. Whether the matched group is best or not is selected relative to other groups. Thus, sub-groups with low *similarity* may be grouped together when there are no better options. It’s reasonable to filter out low *similarity* groups and treat them as unmatched groups for downstream analysis. Refine marker genes list might improve the *similarity* between matched groups.

### Separability

We propose “*separability*” to quantify the difference in matched sub-groups using the expression level of genes after dimension reduction. *Separability* can be defined for the entire transcriptome or the set of highly variable genes from the pre-analysis. *Separability* is defined based on the distance of the K-nearest neighbors intra- and inter- samples in a two-dimensional space including all cells from datasets (2D space such as the combined t-SNE plot, Figure S1D). For each new group and each pair of two samples in the group, the *separability* index is defined as the median difference of intra- and inter-sample distance of each cell within the new group. Assume **C_i_^(1)^** is a cell from sample 1 with n1 cells, we search for the k nearest cells to **C_i_^(1)^** for all cells within sample 1, represented by **C_ik_^(1)^**. We also search for the k nearest cells to **C_i_^(1)^** for all cells within sample 2, represented by **C_ik_^(2)^**, sample 2 with N2 cells. We define

Intra-sample distance for cell **C_i_^(1)^** is

> **D_i_^Intra^ = median_k_ || C_i_^(1)^ − C_ik_^(1)^ ||, for k=1, 2, 3, …, K**
Inter-sample distance for cell **C_i_^(1)^** is

> **D_i_^Inter^ = median_k_ || C_i_^(1)^ − C_k_^(2)^ ||, for k=1, 2, 3, …, K**
The *separability* of sample1 to sample2 is

> **SEP1 = median_i_ (D_i_^Inter^ − D_i_^Intra^), for i=1, 2, 3, …, n1**
For every cell in the second sample, **C_j_^(2)^**, the *separability* of each cell is calculated similarly. The *separability* of sample2 to sample1 is

> **SEP2 = median_j_ (D_j_^Inter^ − D_j_^Intra^), for j=1, 2, 3, …, n2**
Then the *separability* of sample1 vs sample2 in this new group is defined as

> ***Separability* = mean (SEP1, SEP2)**

Using median instead of mean will reduce variation due to outliers. Increasing K will improve accuracy but slow down computation. Practically, the default K is 5, using K > 20 improves the accuracy only slightly.

### Case studies

#### Epithelial cells in different estrus cycle phases

We first applied ClusterMap to compare sub-population changes in two different biological phases using the epithelial datasets that were generated in the study of Pal et al. (15). Cells were collected from mammary glands of adult mice during different phases of the estrus cycle. By pooling the glands from two mice, scRNA-Seq of 2729 epithelial cells in estrus and 2439 cells in diestrus was performed using the 10X Chromium platform.

We analyzed individual datasets using the Seurat package and identified marker genes for 13 sub-groups in diestrus and 10 sub-groups in estrus (Figure 3A, 3B). To match between the set of 13 diestrus groups and the set of 10 estrus groups, ClusterMap clustered the subgroups based on all marker genes identified during pre-analysis (Figure 3C). The clustering dendrogram (Figure 3D) was decomposed using purity tree cut algorithm to form new matched groups (Methods, Figure 3F). Matched sub-groups are shown in Figure 3F and are connected with chords in the Circos plot (Figure 3E). The *similarity* of matched groups ranges from 0.29 to 0.65 (Figure 3F) and is shown in the transparency of the chords in the Circos plot. This indicates that the sub-groups were matched with different percentage of marker genes overlapped. The population size changes were shown by the cell percentage in Figure 3F and indicated by the black sectors in Circos plot. The regroup 2, 3 and 9 were obviously increased in estrus (14% to 24%, 8% to 35%, and 1% to 5% respectively, Figure 3F). ClusterMap recolored the t-SNE plots of each sample (Figure 4A, 4B) and the combined samples (Figure 4C) with the new group assignments. Matched groups are shown in the same color and with the same new group label. Cells with the same color were clustered together in the combined sample, which confirmed our matching results. Additionally, the *separability* for each new group further highlighted the most affected sub-groups. The higher the *separability* value, the more drastically the group was changed, such as regroup 6, 8 and 3, in addition to the groups unique to one sample (Figure3F).

In this case, there are three types of matching sub-groups: one-to-one, one-to-multiple, and singleton. For example, regroup 1 matched p7d_4 and p7e_1, which are tightly clustered together in the combined sample (Figure 4D). Regroup 2 and 3 matched more than one sub-group from the estrus sample to a single group in the diestrus sample. For both cases, we can see cells in these two sub-groups (p7e_3 and p7e_4, p7e_0 and p7e_2) in estrus are closely related and adjacent to each other (Figure 3B). Regroup 7,10, and 11 only include sub-groups from diestrus with no matching sub-groups from estrus, suggesting these sub-groups uniquely exist in diestrus.

Based on known markers, we can label regroup 2, 5, 6, and 10 as mature luminal (ML), regroup 1, 11, and 12 as luminal progenitors (LP), group 9 as luminal intermediate (Lum Int), and regroup 3, 4, 7, and 8 as basal cells (Figure S2A-B, 4A-D). In line with Pal’s observations, we note that the basal population increases slightly in estrus (43% to 47%), ML becomes two major sub-types (regroup 2, 10) and the Lum Int is substantially reduced (5% to 1%) in the diestrus phase. However, ClusterMap unveils more detailed changes between diestrus and estrus that were not identified previously. We found that in basal, regroup 3 was increased substantially in estrus (8% to 35%) with large *separability* (2.93), while regroup 7 is missing in estrus (Figure 3F, 4A-D). Also, regroup 8 in basal is substantially altered between the two phases with a *separability* of 7.13. In the ML population, we noticed that although both group 2 and 10 are ML, regroup 2 in diestrus resembles ML in estrus more closely than regroup 10 (Figure 3D, F). While cells in regroup 2 are a mixture of both phases, cells in regroup 10 are exclusively from the diestrus phase (Figure 4C, 4D). These suggest that a subset of ML cells in diestrus begin to diverge, but ML cells in estrus are more homogenous. Pal et. al also observed that one of the ML subtypes was “tightly associated with ML signature genes such as PgR”, but they neglected to identify that there were different relationships between the two ML subtypes in diestrus and the ML in estrus. One of the ML subtypes in diestrus is much closer to the ML in estrus. For the LP population (regroup 1, 11 and 12), our analysis suggests regroup 11 and 12 are unique sub-types in diestrus (Figure 4A-D), while Pal et. al’s analysis concluded that the LP population was unaltered.

We further characterized the unique groups (7,10, and 11) in the diestrus phase through gene ontology and pathway analysis (21). We found that the marker genes of these groups were extremely enriched for the terms of *ribosome biogenesis, oxidative phosphorylation* and *metabolic process of ribonucleotides* (Figure 4E-F, S2C-D). These findings reflect the increased levels of progesterone in diestrus, which functions as a potent mitogen to stimulate expansion of mammary epithelia at this stage. Only some subpopulations of basal, ML and LP in diestrus responded to progesterone to undergo rapid cell cycle progression, potentially suggesting the existence of differential regulation of mitogen-related cell signaling among sub-populations of the same cell type. Intriguingly, we noted group 10 cells in diestrus are associated with the development of mammary gland alveoli, indicating that some of the cells within this group possess trans-differentiation potential for alveolar development (Figure 4F)(22). In addition, we observed a drastic increase in the number of group 3 cells with characteristics of smooth muscle in estrus, indicating there may be a contractile switch of myoepithelial cells for lactation preparation during this period (Figure S2D).

For comparison with ClusterMap, we also performed canonical-correlation analysis (13) on the epithelial cell datasets (Figure 4G, S5). The two samples mixed more evenly under canonical correlation vector space (Figure S5B). Note that the two separated groups, group 2 and group 10 (Figure 4A) of mature luminal (marked by Prlr) in diestrus were not separated into sub-groups after the CCA analysis (Figure 4G, S5A-B). This indicated that CCA reduced the difference between sub-groups even in the same sample, which led to existing sub-groups becoming less distinguishable. Butler et al. observed this effect as well for rare populations and suggested using PCA for further analysis (13). However, the sub-group 10 in the diestrus phase (Figure 4A, 4C) was not a rare population. With CCA only, we may miss the detection of these sample specific sub-groups identified by ClusterMap. Thus, CCA and ClusterMap provide different views of comparing multiple scRNA-Seq datasets.

#### PBMCs under immune stimulus

We next applied ClusterMap to compare effects across experimental treatments on each cellular population. The datasets we used were generated in the study of Kang HM, et al. (16). Peripheral blood mononuclear cells (PBMCs) from each of eight patients were either untreated as a control or activated with recombinant interferon-beta IFN-β for 6 hours. The same number of IFN-β-treated and control cells from each patient was pooled and subjected to single-cell sequencing on a 10X Chromium instrument. Transcriptomes of 14,619 control and 14,446 IFN-β-treated single cells were obtained.

The pre-analysis using Seurat defined 11 and 13 sub-groups for the control and stimulated conditions respectively (Figure S3A-B). With hierarchical clustering and the purity tree cut approach (Figure 5A-B), ClusterMap matched most sub-groups, except the regroups 10, 11 and 12 (Figure 5C, S3B). We confirmed that the matched groups expressed the same known marker genes, demonstrating that matching worked correctly (Figure S3C-D). There is no obvious change in the percentage of each cell type after stimulation, which is consistent with Kang et al.’s conclusion. For group 10 and 12, although both groups express some megakaryocyte marker genes, such as PPBP, other marker genes of group 10 and 12 do not overlap with each other very well, placing them in distantly related clusters (Figure 5A arrows). Thus, ClusterMap considered these two sub-groups to be distinct new groups. However, group 10 and 12 partially overlap in the combined analysis (Figure 5F). We suspect that this inconsistency might be due to the relatively small population size in both sub-groups. The small population size increases the false positive rate of identified marker genes, which negatively affects the performance of ClusterMap.

**Figure 5.**
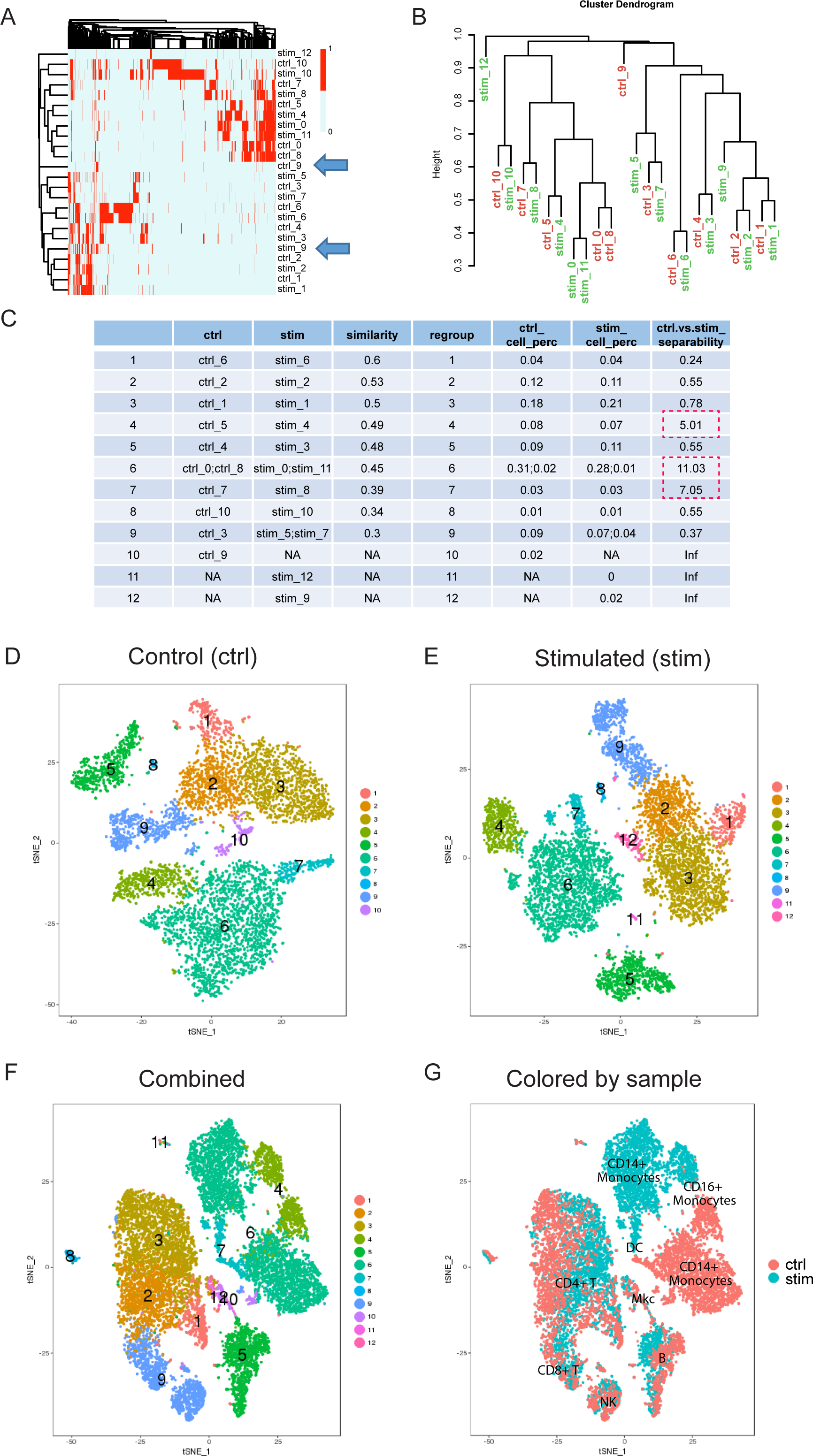
ClusterMap analysis for the immune stimulated datasets. A. Heat map of marker genes. B. Dendrogram of the hierarchical clustering of marker genes. C. ClusterMap results for the quantification of the sample comparisons. D-E. Re-colored t-SNE plot based on matching results for control and stimulated conditions. F. Re-colored t-SNE plot based on matching results for the combined dataset. G. t-SNE plot with cells colored by sample. Based on known marker genes, we assigned group 1, 2, 3 to CD4+ T cells, group 5 to B cells, group 6 to CD14+CD16-monocytes, group 4 to CD14+CD16+ monocytes, group 7, 8 to dendritic cells, group 10, 12 to megakaryocytes, group 11 to erythroblast and group 9 to a mixture of natural killer cells and CD8+ T cells (Figure S3C-D).

We observe a wide range of *separability* measures across matched sub-clusters, indicating the different sub-groups responsd to IFN-β stimulation very differently. For instance, T cell group 1 was less affected, while the IFN-β stimulation affected monocytes and dendritic cells much more than the other immune cells (Figure 5C red dashed boxes). This is confirmed in the t-SNE plot from the combined analysis, the distance of cells from the two conditions in group 4, 6, and 7 are much farther than the other groups (Figure 5F-G). In addition, we note that although regroup 5 and 6 match with comparable *similarity* (0.48 and 0.45), the *separability* of the two groups is quite different (0.55 and 11.03). This suggests that the overlapping of marker genes is similar for the two regroups, but the changes in transcriptome expression levels may be drastically different.

Butler et al. applied CCA to the same immune stimulated dataset (13). By aligning each cell, the samples are scaled to become as similar as possible. The global differences in the original datasets were treated as a batch effect instead of biological. However, the IFN-β stimulus was expected to trigger a widespread immune response of PBMCs (16). It is hard to determine if the effect is due to biology or technique. Additionaly, we observed dendritic cells (DC, regroup 7) and monocytes (regroup 4, 6) respond to IFN-β the most, while Butler observed plasmacytoid DC respond to the stimulus the most, but did not see an obvious response for the myeloid and lymphoid cells.

#### PBMCs replicates as a control

To determine if our method introduced any spurious matches, we used ClusterMap to compare two replicated datasets. Ideally, clusters in replicated data should match each other in a one-to-one manner with minimal differences observed. The 4K PBMC and 8K PBMC datasets were downloaded from the 10X Genomics public datasets (https://support.10xgenomics.com/single-cell-gene-expression/datasets/). They are peripheral blood mononuclear cells (PBMCs) from the same healthy donor. The two datasets are replicates with different detected cell numbers, with about 4000 and 8000 cells each.

The group matching results demonstrate that the sub-groups in the 4K PBMCs match with the 8K PBMCs in a one-to-one manner (Figure 6). The *similarity* values between matched groups is also much higher than in the other two datasets (Figure S4D). The chord color in the circos plot is hence much darker (Figure 6B). The majority of the cell percentages of sub-groups are not changed. The *separability* values demonstrate that there are no obvious differences between the two samples in any of the matched pairs (Figure S4D). This matches the expectation as 4K and 8K PBMCs are replicates and shows that ClusterMap will not find spurious relationships where they don’t exist in the data.

**Figure 6.**
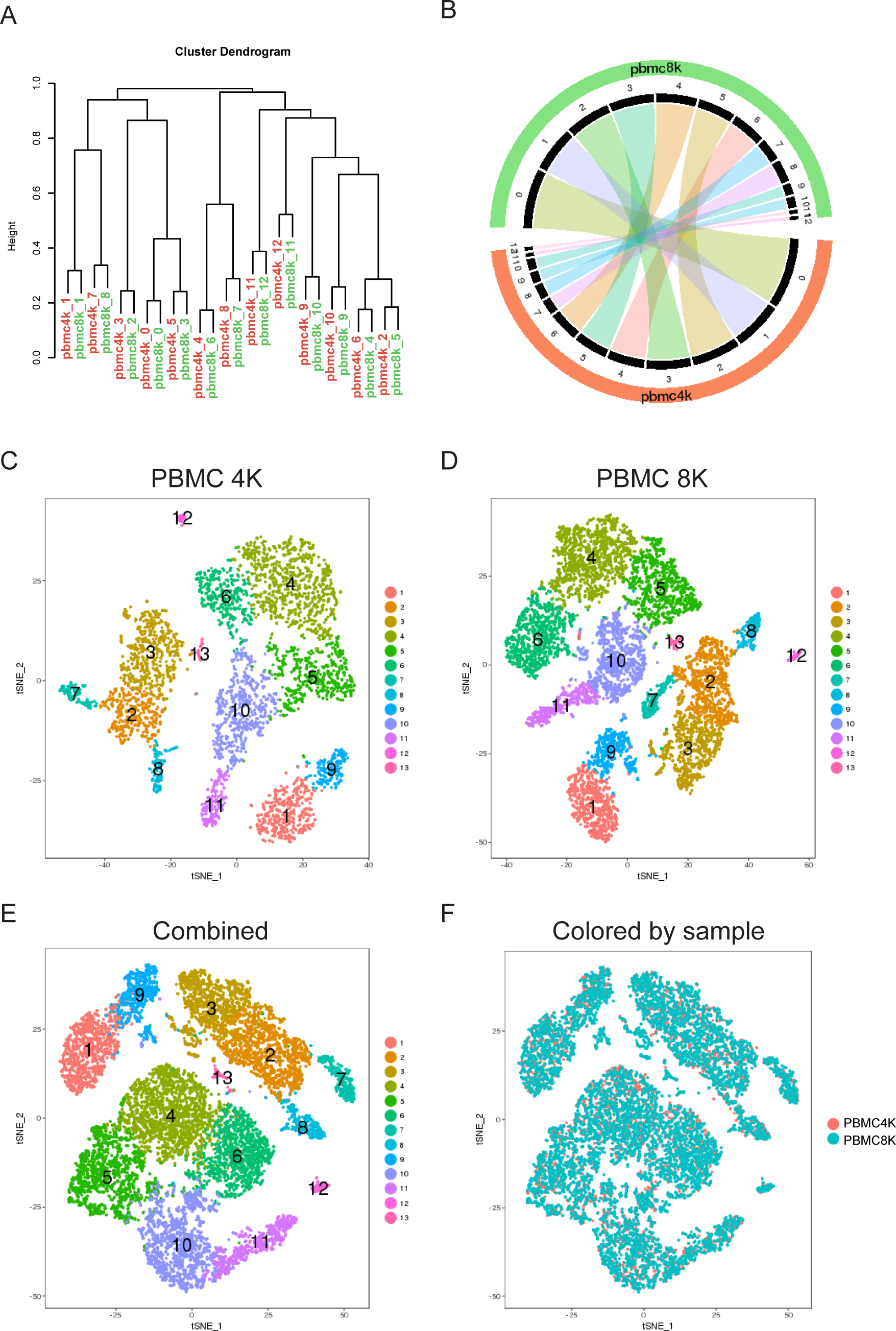
ClusterMap analysis for the PBMC replicates datasets. A. Dendrogram of the hierarchical clustering of marker genes. B. Circos plot of the matched subgroups. C-D. Re-colored t-SNE plot based on matching results for control and stimulated conditions. E. Re-colored t-SNE plot based on matching results for the combined dataset. F. t-SNE plot with cells colored by sample.

## Discussion

Due to the limitation of current scRNA-seq experimental design, batch effects and biological effects are always confounded. It is challenging to distinguish batch effect from sample differences. During the matching step of ClusterMap, batch effects will not affect the matching results. Because marker genes for each sub-group are identified relative to the rest of the sub-groups within a given sample, matching based on the existence (binary) of marker genes will overcome the batch effect. *Separability* can indicate the existence of batch effects or systematic variation, if large *separability* values are observed over all matched groups. There are many studies on comparison of multiple scRNAseq datasets from different batches, such as the mutual nearest neighbors method (MNNs, (11)) or the distribution-matching residual networks method (12). If necessary, *separability* can be applied after batch effect correction.

As a downstream analysis approach of scRNA-seq, ClusterMap relies on the pre-analyzed data. The clustering analysis of each single sample and the marker genes identified for each sub-group will affect the quality of the matching results. Refining marker gene lists will certainly improve the sub-group matching. It’s important to define meaningful sub-groups for each sample first before starting a cluster comparison. The regroup step in ClusterMap refines the clustering based on the matched results, possibly merging some similar sub-groups in the same sample.

*Similarity* measures how well a pair matches each other compared to other groups based on the marker genes. *Separability* measures how much the group properties change between paired groups. They can both reflect the relationship of the paired groups, but in different ways. Typically, as *similarity* rises, *separability* decreases. However, it is possible that a pair has similar marker genes, but separates far apart (group 6, Figure5C, F, G). This is due to the use of binary values of marker genes to compare between groups while expression level is used to compare cell distance within paired groups.

The clustering analysis may be performed on the combined samples as well. Better resolution will be gained by increasing the cell numbers of the pooled dataset. However, the new clustering results will be hard to match back to the sub-groups in each single sample. The regrouping results in ClusterMap keep the grouping information for the single samples. It also makes sense to compare the match-defined new groups and the clustering-analysis-defined new groups in the combined sample.

We developed CluserMap to systematically compare multiple scRNA-seq datasets at the cluster level. We objectively match sub-groups using marker genes and purity tree cut methods. The *similarity* quantifies the matching quality and the *separability* characterizes the changes within the matched groups. Unlike CCA and scMAP, we are not trying to align cells across datasets, rather, we focus on how sub-groups change. Our case studies demonstrate that Clustermap facilitates the detection of sample specific cell types and cell types that change with experimental conditions. By providing clear visualization of the results with multiple customized plots, ClusterMap enables us to gain more insight into biological mechanisms at the sub-population level. In comparison with other existing methods, we provide a higher-level approach to comparing multiple scRNA-seq samples, with more quantifications and visualizations to allow for easier interpretation and downstream analysis.

## Availability

An R package and documentation for ClusterMap is deposited in a GitHub repository (https://github.com/xgaoo/ClusterMap). The code and results for the pre-analysis of the datasets in this study are also available.

The epithelial datasets and immune stimulated datasets were downloaded from Gene Expression Omnibus under accession numbers GSE103272 and GSE96583. The PBMC replicates datasets were downloaded from the 10X Genomics support datasets website at (https://support.10xgenomics.com/single-cell-gene-expression/datasets/2.1.0/pbmc4k and https://support.10xgenomics.com/single-cell-gene-expression/datasets/2.1.0/pbmc8k).

